# Interictal epileptiform discharges affect memory in an Alzheimer’s Disease mouse model

**DOI:** 10.1101/2023.02.15.528683

**Authors:** Marisol Soula, Anna Maslarova, Ryan E Harvey, Manuel Valero, Sebastian Brandner, Hajo Hamer, Antonio Fernández-Ruiz, György Buzsáki

## Abstract

Interictal epileptiform discharges (IEDs) are transient abnormal electrophysiological events commonly observed in epilepsy patients but are also present in other neurological disease, such as Alzheimer’s Disease (AD). Understanding the role IEDs have on the hippocampal circuit is important for our understanding of the cognitive deficits seen in epilepsy and AD. We characterize and compare the IEDs of human epilepsy patients from microwire hippocampal recording with those of AD transgenic mice with implanted multi-layer hippocampal silicon probes. Both the local field potential features and firing patterns of pyramidal cells and interneurons were similar in mouse and human. We found that as IEDs emerged from the CA3-1 circuits, they recruited pyramidal cells and silenced interneurons, followed by post-IED suppression. IEDs suppressed the incidence and altered the properties of physiological sharp-wave ripples (SPW-Rs), altered their physiological properties, and interfered with the replay of place field sequences in a maze. In addition, IEDs in AD mice inversely correlated with daily memory performance. Together, our work implicates that IEDs may present a common and epilepsy-independent phenomenon in neurodegenerative diseases that perturbs hippocampal-cortical communication and interferes with memory.

**Significant Statement:** Prevalence of neurodegenerative diseases and the number of people with dementia is increasing steadily. Therefore, novel treatment strategies for learning and memory disorders are urgently necessary. IEDs, apart from being a surrogate for epileptic brain regions, have also been linked to cognitive decline. Here we report that IEDs in human epilepsy patients and AD mouse models have similar local field potential characteristics and associated firing patterns of pyramidal cells and interneurons. Mice with more IEDs displayed fewer hippocampal SPW-Rs, poorer replay of spatial trajectories, and decreased memory performance. IED suppression is an unexplored target to treat cognitive dysfunction in neurodegenerative diseases.

## Introduction

The hippocampus is characterized by the largest recurrent excitatory network in the mammalian brain (Amaral & Witter 1989; Li et al., 1994; Wittner et al. 2007). In the intact brain, two characteristic local field potential (LFP) patterns, theta oscillations and sharp-wave ripples (SPW-Rs) are associated with appetitive and consummatory classes of behavior, respectively (Buzsáki 2015). SPW-Rs arise from the synchronous discharges of CA3 pyramidal cells via their spatially extensive excitation. The CA3 population discharge gives rise to the LFP sharp wave (SPW) in the target apical dendrites of CA1 neurons in the str. radiatum (Buzsáki et al., 1983). In turn, the strong excitation of CA1 pyramidal cells and interneurons induces a short-lived and fast (40-150 ms; 100-150 Hz) oscillation, known as the ripple (Buzsáki et al. 1992; Stark et al., 2014). The physiological significance of SPW-Rs is supported by numerous observations in humans and experimental animals, demonstrating that the neuronal discharge patterns of learning are replayed in a compressed manner during SPW-Rs (Wilson and McNaughton, 1994; Vaz et al., 2020) and that aborting and prolongation of SPW-Rs reduce and enhance memory, respectively (Girardeau et al., 2009; Jadhav et al., 2012; Fernández-Ruiz et al., 2019).

Alteration of hippocampal circuits and their physiological dynamics is believed to underlie several diseases, including epilepsy and Alzheimer’s disease (AD; Buzsáki & Watson 2012; Staba et al., 2002; Chauvière et al., 2009; Palop & Mucke 2016; Jelic et al., 2000; Jeong 2004; Herrmann & Demiralp 2005; Engels et al., 2016; Allen et al., 2007). In epileptic patients, hypersynchronous events often occur between the relatively rare seizures, known as interictal epileptiform discharges (IEDs; Alarcon et al., 1997; de Curtis et al., 2012). IEDs are assumed to be the result of seizures, and, in turn, their occurrence has been hypothesized to play a role in seizure induction (Staley et al., 2011; Gotman 1991; Gotman & Marciani 1985). Early experiments have hypothesized that IEDs, at least some of their forms, arise from the same CA3 recurrent circuit as SPW-Rs (Behrens et. al., 2007; Lopes da Silva et al., 1995; Buzsáki et al., 1991). In support of this hypothesis, IEDs and SPW-Rs in epileptic rodents and patients compete and IEDs are associated with decreased memory performance (Gelinas et al., 2016; Henin et al., 2021).

IEDs have also been detected in patients with AD and frontotemporal dementia as well as in AD rodent models (Lam et al., 2017; Horváth et al., 2017; Vossel et al., 2016; Kam et al., 2016; Bezzina et al., 2015; Reyes-Marin et al., 2017). People with epilepsy develop AD at a rate several times higher than non-epileptic individuals (Hauser et al., 1986; Amatniek et al., 2006; Pandis et al., 2012). These pathophysiological patterns have been assumed to contribute to cognitive decline and disease progression (Aldenkamp & Arend 2004; Kam et al., 2016; Lam et al., 2017; Vossel et al., 2017; Verret et al., 2012; Volicer et al., 1995). However, AD patients and AD rodent models rarely display epileptic seizures, therefore it is unlikely that IEDs in degenerative disease are brought about by seizures per se. In fact, IEDs appear prior to the histopathologically detected neuronal degeneration in several AD mouse models (Busche et al., 2012; Kam et al., 2016; Reyes-Marin et al., 2017; Gureviciene et al., 2019). Thus, the relationship between bona fide IEDs in epilepsy and the pathophysiological patterns in AD models needs to be clarified. Furthermore, the progression of interaction between IEDs and SPW-Rs and their relationship to memory impairment in AD models awaits clarification. To shed light on these issues, we compared IEDs in human patients with IEDs in AD mice, followed the progression of both IEDs and SPW-Rs for several months, and show that IED prevalence in aged AD mice correlates with memory impairment.

## Results

### IED features in drug-resistant epileptic patients with intracranial LFP

We first investigated the LFP properties of human interictal epileptiform discharges recorded from mesial temporal lobe structures (anterior hippocampus and parahippocampal regions, see Supplementary Table 1) of patients with drug-resistant frontal and temporal lobe epilepsy. We analyzed signals from 7 afternoon and nighttime sessions (median recording time per session 16.54 hours) from 6 patients. All patients were implanted with one to two micro-electrode bundles in the anterior hippocampus, in addition to standard depth macroelectrodes, (Fig. 1A). The combined electrode configuration included 8 macro contacts located in the neocortex and a bundle of 8 microwires targeting the hippocampus (Misra et al., 2014; Fried et al., 1999) Macro/ Behnke-Fried Micro depth electrodes, Ad-Tech Medical, USA). Locations of the macroelectrodes and microwire bundles were confirmed with MRI and CT (Yang et al., 2012; Ekstrom et al.,2008; Misra et al., 2014). For the purpose of this study, we analyzed IEDs recorded from the microwires in the hippocampus (Fig. 1B). The waveforms of IEDs detected on the microwires varied both in amplitude and complexity (Fig. 1B). Because the locations of the individual microwire tips could not be verified with current methods, the waveforms shown in Fig. 1B were rendered heuristically on the basis of measured depth profiles in rodents (Fig. 2B).

**Figure 1:**
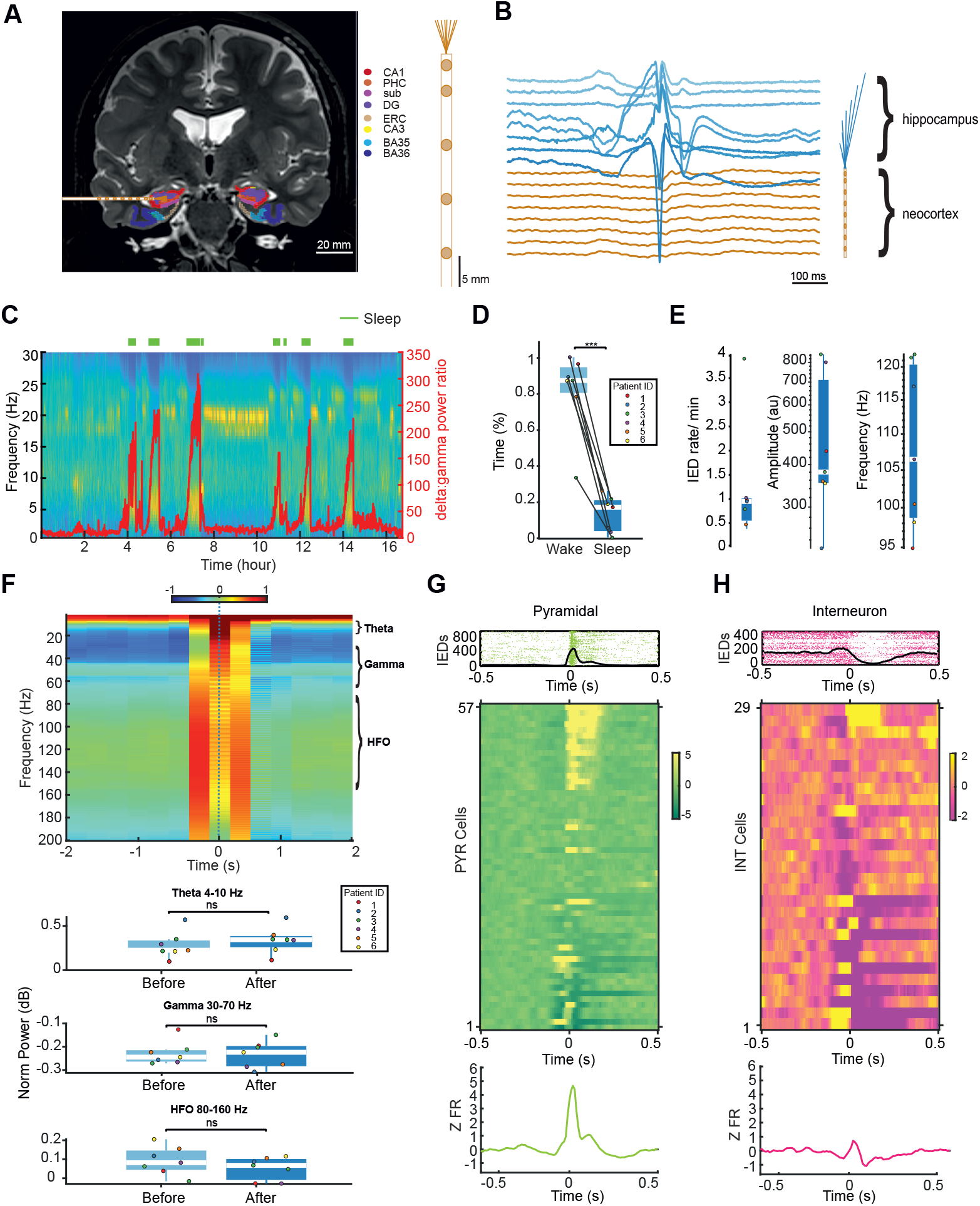
Characterization of IEDs in patients with drug-resistant epilepsy with intracranial EEG (iEEG). **A**. Position of an electrode targeting the hippocampus (Behnke-Fried macro-/microwire electrode) consisting of 8 macro contacts and a bundle of 8 microwires. Mesial temporal structures are color-coded on the T2 MRI scan: red – CA1, orange – perihippocampal cortex, magenta – subiculum (sub), violet – dentate gyrus (DG), tan – entorhinal cortex (ERC), yellow – CA3, light blue – Brodmann area 35 (BA35), dark blue – Brodmann area 36 (BA36) (Yushkevich et al., 2015). Right: schematic of the electrode (the last three outer macro contacts are not shown). **B**. Left: LFP traces of an IED recorded across all channels from one temporal electrode – channels 1-8 are hippocampal microwires, channels 9 -16 are the macro-contacts on the same electrode. **C**. An example of a recording session with the patient’s sleep-wake scoring (green lines, sleep). **D**. Fraction of awake time vs sleep time in n=6 patients (two-sided Wilcoxon rank-sum test; p = 0.0017). **E**. IED rate, peak frequency, and amplitude of all patients (n = 6). Each color corresponds to a patient. **F**. Top: Example of time-frequency spectrogram during all the IEDs in one session, centered on IEDs (time 0). Bottom: Comparison of the frequency spectrogram 2 s before and 2 s after an IED. Average change in theta (4– 10 Hz; p = 0.277), gamma (30–70 Hz; p = 0.749) and higher frequency oscillations (80-160 Hz; p=0.338) power before and after an IED. (ns= not significant; two-sided Wilcoxon rank-sum test). **G**. Top: Example of a pyramidal cell modulated by IEDs. Bottom: Peri-event firing rates around IEDs of all pyramidal (n= 57 cells) and their average response. **H**. Top: Example of interneurons modulated by IEDs. Bottom: Peri-event firing rates around IEDs of all interneurons (n= 29) and their average response. Color axis is z-scored firing rate [-5 5] or [-2 2].

**Figure 2:**
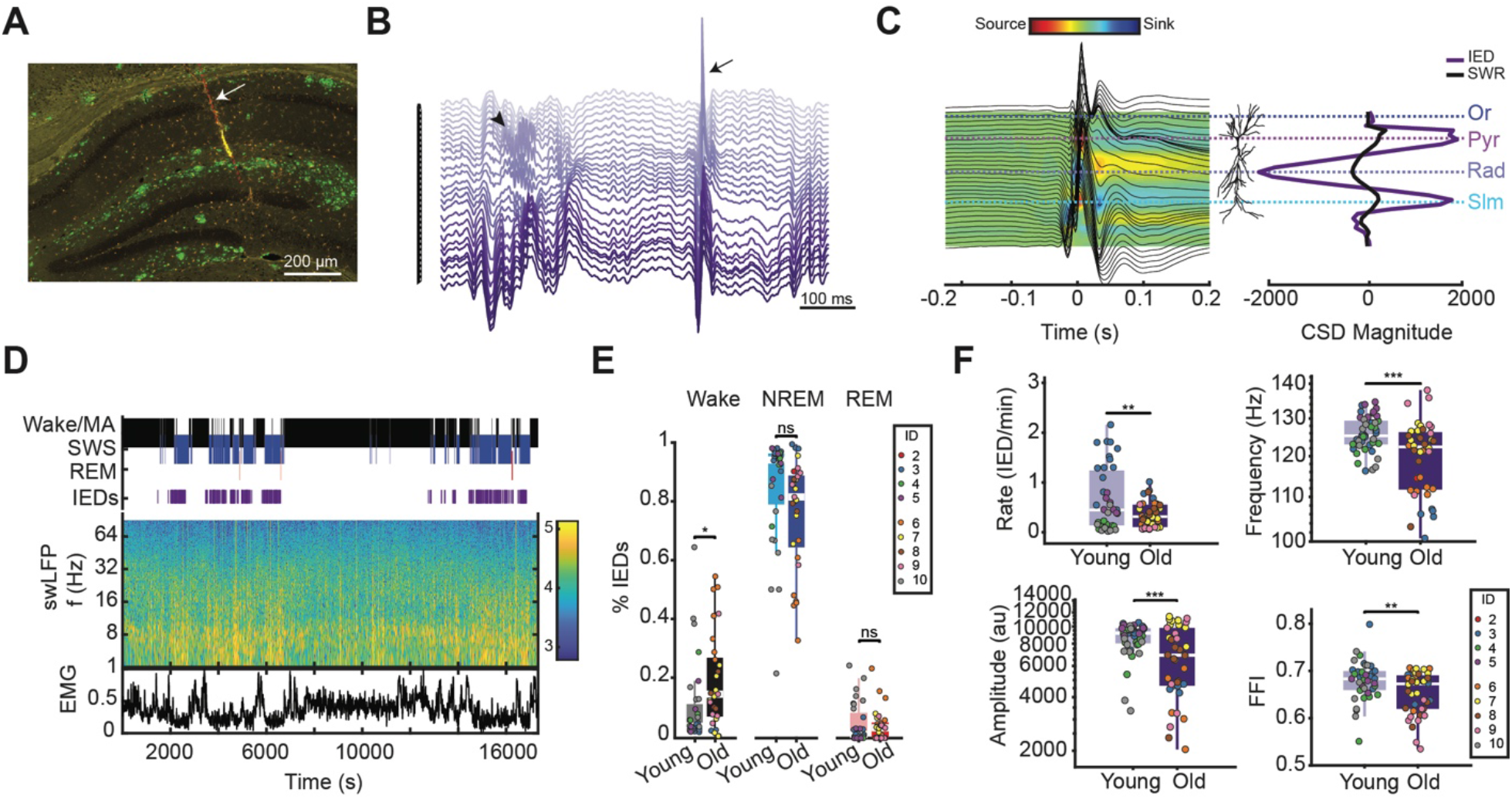
Characterization of IEDs in APP/PS1 AD mouse model. **A**. Arrow points to track of one shank of the two-shank, 64-site linear silicon probe (red line) spanning the CA1-dentate axis. Green= D54D2; anti-β-Amyloid antibody. Red= IBA-1; anti-microglia). **B**. LFP traces of SPW-R (arrowhead) and an IED (arrow), recorded across all channels from one shank. **C**. Left: LFP traces overlayed on the CSD map of an IED. Horizontal dashed lines correspond to the str. Oriens (Or), str. pyramidale (Pyr), str. radiatum (Rad), and str. lacunosum-moleculare (Slm). Right: depth profile of CSD sinks and sources at the peaks of averaged IEDs and SPW-Rs. **D**. Example recording session with state scoring, EMG, and occurrence of IEDs. **E**. Group data for IED occurrence during wake, nonREM and REM in young (n=4) and old mice (n=6) (two-sided Wilcoxon rank-sum test, p = 0.0097, p=0.0108, p=0.636; ns = not significant). **F**. Group differences of IED rate, frequency, amplitude, and fast frequency index in young (n=4) and old mice (n=6) (two-sided Wilcoxon rank-sum test, p = 0.1415; p=0.0042; p=0.0012; p=0.014). Each dot corresponds to a given session and the color to an animal.

Implanted patients were resting in a hospital bed with limited mobility during the recording period. Patients slept irregularly, sleeping in bouts of 1 hr at a time (Fig. 1C). Although recordings included late afternoon and early night, the patients were mostly awake during the recording epochs (0.87 ±0.09% wake; Fig. 1D). NREM sleep facilitates the production of interictal spikes in medial temporal regions (Lambert et al., 2018). For detecting IEDs, the wide band signal was band-pass filtered (20-80 Hz) and transformed to a normalized squared signal (NSS). IEDs were detected as peaks beyond 10 standard deviations (SD) of the mean NSS and when the signal crossed 3 SDs of the mean NSS for 30 - 250 ms. Throughout the recording sessions, IEDs occurred at a rate of (1.23± 1.21 /min) and the average peak amplitude was 485.83± 239.06 µV. The peak frequency of the IEDs, as determined from the power spectrum of the detected IEDs, was 108.63±11.42 Hz. To uncover the relationship of IEDs with the on ongoing brain state, we compared the spectral components of the LFP signal 2 s before and after the IEDs (Fig. 1F). Although comparison of the spectral bands before and after the IEDs in the theta (4-10 Hz), gamma (30-70 Hz) and high-frequency oscillations (80 – 160 Hz) bands showed no significant difference (theta p = 0.277, gamma p = 0.749, HFO p=0.338), likely due to the small number of sessions, the direction of changes (increased gamma power and decreased 80 – 160 Hz band power) was similar to those reported previously (Wu et al., 2021; Meisenhelter et al., 2021).

In addition to LFP, the microwires also detected neuronal spikes. After spike clustering, we classified the recorded units into 57 putative pyramidal cells and 29 interneurons according to firing rate and spike waveforms features (Supplementary Fig. 1A-C), and analyzed their firing modulation before, during and after IEDs. We found that 47% of putative pyramidal cells and 38% of putative interneurons were modulated by IEDs (Supplementary Fig. 1D). IEDs were accompanied by a synchronous increase of pyramidal neuron firing rate followed by a transient suppression of spiking. Interneurons showed a heterogeneous response from increased or suppressed spiking during the IED followed by quick recovery or suppression (Fig. 1G). Using these characteristics of IEDs in humans, next we compared them with similar features of IEDs in AD mice.

### IEDs in AD mouse model

Previous studies have already reported IEDs and epileptic seizures in APP/PS1 mice on a C57BL/6J genetic background (Reyes-Marin & Nuñez 2017; Gureviciene et al., 2019; Minkeviciene et al., 2009). Mice used in this study, APP/PS1 hemizygotes on a C57BL/6;C3H-congenetic background (JAX Stock No. 004462), have alterations in excitatory/inhibitory (E/I) balance but no known report of IEDs has been documented (Algamal et al., 2021; Algamal et al., 2022; Yang & Jeong 2021). Mice were implanted with a two-shank linear silicon probe in the hippocampus spanning the CA1 layers and the upper blade of the dentate gyrus (Fig. 2A). IEDs were detected in every APP/PS1 mouse at all ages (n=10), spanning from 2 to 13 months but not in control mice (n=6). These short and complex waveform events were strikingly different from the background activity and SPW-Rs (Fig. 2B). To study their anatomical layer characteristics, we constructed peak-triggered laminar averages of the LFPs and the corresponding current-source density (CSD) maps and compared them with the laminar distribution of LFP and CSD profiles of SPW-Rs. The depth profiles of IEDs and SPW-Rs were remarkably similar, although the magnitude of IEDs were several times larger than that of SPW-Rs (Fig. 2C).

The majority of IEDs occurred during NREM sleep (Fig. 2D-E) and often in bursts (Supplementary Fig. 2D). In old mice the occurrence IEDs was higher during waking compared to young mice (Fig. 2E). In all recording sessions, the overall mean IED rate was significantly lower in old compared to young mice (Fig. 2F; young: 0.67±0.65/min, old: 0.32±0.26/min). Both the peak amplitude and peak frequency of the IEDs were also reduced in old compared to young animals (Fig. 2F; Supplementary Fig. 2A-C). We also computed a “fast frequency index”, a measure of the fraction of power above 250 Hz in the 100–600 Hz spectral band (Valero et al., 2017), which also decreased significantly in old mice (Fig. 2F; young: 0.69 young; old: 0.66).

In addition to animal group comparisons, we performed longitudinal recordings of physiological event as young mice grew older. In two AD mice and two littermate control mice, we monitored electrophysiological activity and behavior weekly from 2.5 months of age to 13 months. In AD mice the rate of IEDs decreased over time, spatial memory performance decreased, the firing of pyramidal cells increased, and interneurons decreased, SPW-R power decreased over time, while the incidence of SPW-R remained constant. Control mice showed no significant or less pronounced changes in these electrophysiological markers during the same period (Supplementary Fig. 3D; Supplementary Fig. 7A-C).

### IEDs in AD mouse model alter hippocampal circuit dynamics

Similar to the human recordings, we analyzed the spectral components of the signal 2 s before and 2 s after the IEDs (Fig. 3A). In young mice, theta (4-10 Hz) and gamma power (30-70 Hz) increased following an IEDs. While power in the high frequency oscillation band (HFO; 80-160 Hz) decreased, it was not significant (Fig. 3B). In aged mice, in contrast, the IED-induced decrease of HFO band power was significant, gamma power also increased but to a smaller extent than in young mice, whereas theta band power was not affected (Fig. 3B).

**Figure 3:**
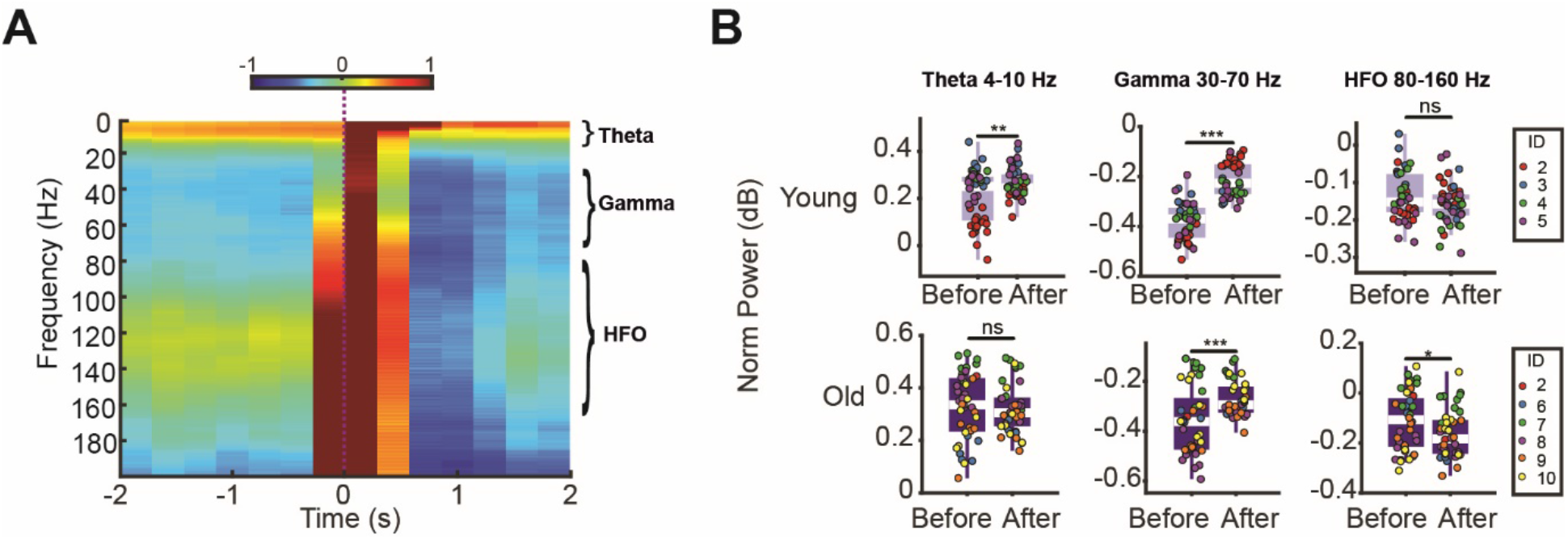
IEDs transiently alter hippocampal LFP. **A**. Example of time-frequency spectrogram centered around IEDs in an example session. Color axis is normalized power [-1 1]. **B**. Average power change in theta (4-10 Hz; two-sided Wilcoxon rank-sum test, p=0.0036, p=0.554), gamma (30-70 Hz; two-sided Wilcoxon rank-sum test, p=0.0007, p=0.00043) and high frequency (80-160 Hz; two-sided Wilcoxon rank-sum test, p=0.227, p=0.0439) bands before and after IEDs in young and old mice. Each dot corresponds to given session and the color to an animal.

To examine the neuronal correlates of IEDs, we separated pyramidal cells and interneurons by their spike duration and bursting index (Petersen et al., 2021; Soula et al., 2023; Supplementary Fig. 3A-C) and quantified their time courses of spiking around IEDs both at short (±0.5s) and longer (±20s) time windows by constructing peri-event time histograms of individual neurons, centered on IEDs. Both pyramidal cells (Fig. 4A) and putative interneurons (Fig. 4B) were suppressed for ∼50-200 ms after an IED but, on average, the post-IED suppression was less pronounced in aged AD mice (Fig. 4C). This suppression was positively correlated with the IED CSD magnitude (Supplementary Fig. 4). The 20 ms time window (± 10 ms) surrounding the peak of the detected IEDs were excluded from this analysis because during the LFP spike component of the IED, many neurons became synchronized and the superimposed spike waveforms prohibited their individual detection. The peak time of population synchrony was comparable between young and aged mice.

**Figure 4:**
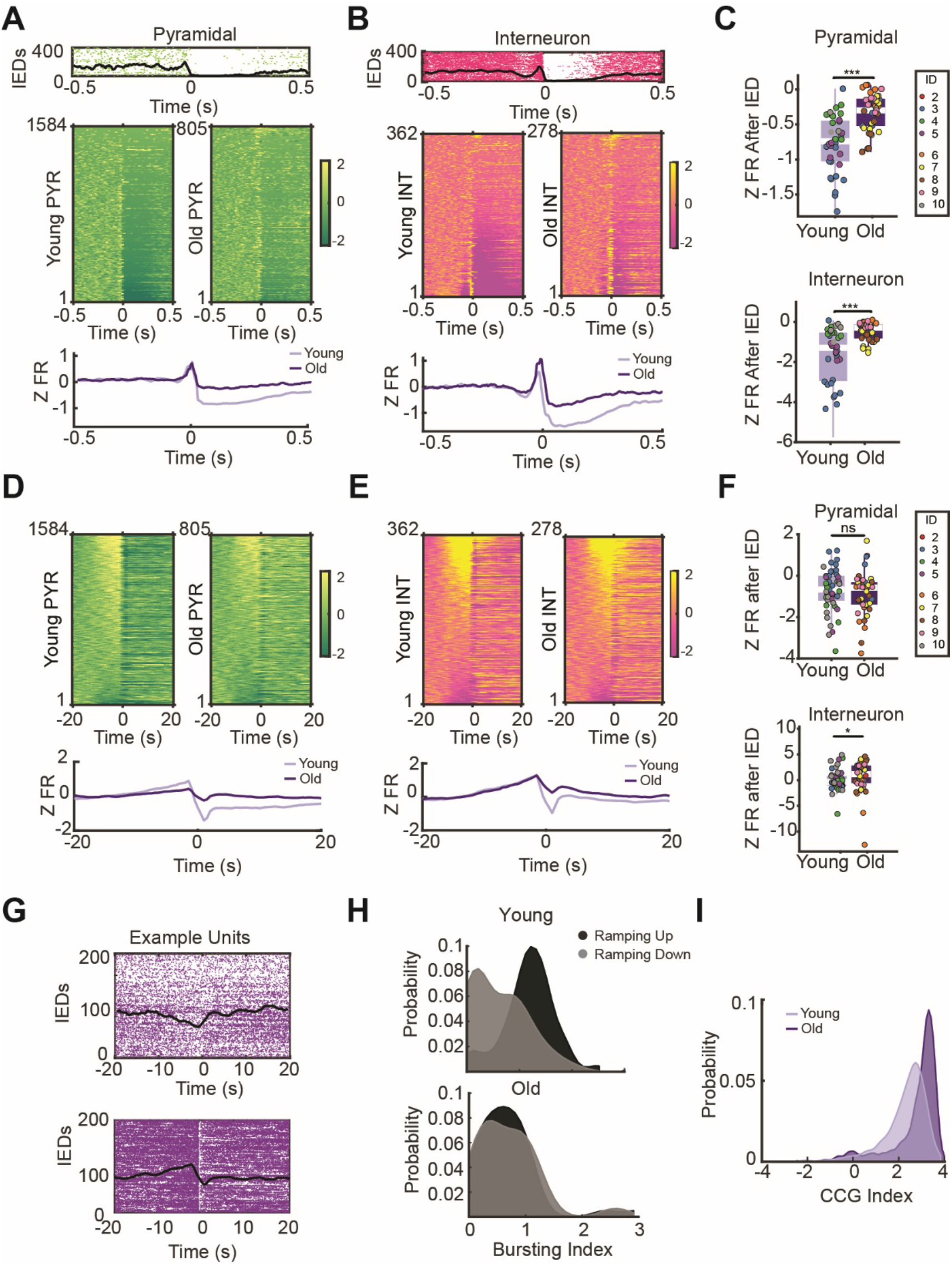
IEDs alter neural spiking at short and long-time scales. **A**. Top panel. Peri-IED example of spiking in a pyramidal cell. Middle panel. Z-scored, peri-IED spiking in all recorded pyramidal cells in young and old AD mice, sorted by strength of post-IED inhibition (bottom, most suppressed neurons). Bottom panel. Mean z-scored firing rates of all pyramidal cells around IEDs. **B**. Same as for putative interneurons. Color axis is z-scored firing rates [-2 2]. **C**. Suppression of z-scored firing rates of pyramidal neurons (top: two-sided Wilcoxon rank-sum test, p = 1.67e-80) and interneurons (bottom: two-sided Wilcoxon rank-sum test, p = 1.30e-6).) in the 0.05-0.5s time window after IED in young and old mice; sided Wilcoxon rank-sum test. Each color corresponds to a given animal. **D**. Mean z-scored firing rates of all pyramidal cells 20 seconds before and after IEDs. All young (left) and old (right) pyramidal cells peri-event z-scored firing rate response 20 seconds before and after an IED. Bottom panel: Average z-scored responses of all pyramidal cells in young (light purple) and old mice (dark purple). **E**. Same as D for interneurons. Color axis is Z scored firing rates [-2 2]. **F**. Differences of z-scored firing rates in the 0.05-5s time window after IED in young and old mice for pyramidal cells (top: two-sided Wilcoxon rank-sum test, p = 0.38) and interneurons (bottom: two-sided Wilcoxon rank-sum test, p = 0.024). **G**. Example neurons that ramp down (top) and ramp up (up) firing rates before IEDs. **H**. Interneurons were separated into ramping up and down cells, and the probability distribution of their bursting index in young (top) and old (bottom) (Royer et al., 2012) is shown for the two groups. (Young: ramping up-mean= 1.1834 std= 0.5009; ramping down-mean= 0.6117 std=0.5812; KW test, p=9.57e-10. Old: ramping up-mean= 0.6758 std= 0.6080; ramping down-mean= 0.6888 std=0.6027; KW test, p=0.941. **I**. Cross-correlogram (CCG) index of all cells in young (mean= 2.7428; std= 1.0836) and old mice (mean= 2.2942; std= 0.9577; KW test, p=9.48e-83).

The transient post-IED suppression of spiking activity was followed by a longer-lasting steady decrease and a ramping up of the firing rate prior to the occurrence of IED of both pyramidal cells and interneurons (Fig. 4D-E). Similar to the transient suppression, the post-IED steady state decrease of interneuron firing was significantly stronger in young compared to aged AD mice (Fig. 4F). In old mice, neuron pairs had a lower cross-correlograms (CCG) index, indicating reduced synchrony. The CCG index was calculated by taking the median z-firing cofiring rate 0.1 ms around the CCG of all cells (Fig. 4I). To further analyze potential age-dependent differences, we calculated the burst index, defined by the fraction of spikes with a neighboring interspike interval less than 6 ms. Interneurons were classified each neuron depending on whether their firing rates prior to the IED increased (“ramping up”) or decreased (“ramping down”). While ramping up and down cells co-segregated with the burst index in young mice, no such segregation was evident in old mice (Fig. 4G-H).

### Relationship between IEDs and SPW-Rs

The results thus far indicate that IEDs are correlated with firing pattern changes of both pyramidal cells and interneurons and that these patterns are different in young and old AD mice. To study the excitability changes further, we examined SPW-R rates during NREM sleep and the relationship between IEDs and SPW-R. Overall SPW-R rate was significantly lower in aged compared to young mice (Fig. 5B; Supplementary Fig. 7). In line with the IED-induced spectral changes and firing rate changes of neurons, the incidence of SPW-Rs preceding and following IEDs showed a characteristic pattern. In young animals, SPW-R rate started to increase steadily at least 10s prior to the IED, followed by a large suppression for ∼2-4 s. Both the pre-IED increase and the post-IED decrease of SPW-Rs were significantly reduced in the old mice (Fig. 5A-B). Post-IED SPW-Rs were associated with significantly reduced firing rate of pyramidal neurons in both young and old mice. In young, but not in old, mice the firing rate of interneurons was also reduced during post-IED SPW-Rs (Fig. 5C). The amplitude, ripple frequency, duration, and entropy features of SPW-Rs did not differ before and after IEDs in either young or old AD mice (Supplementary Fig. 5).

**Figure 5:**
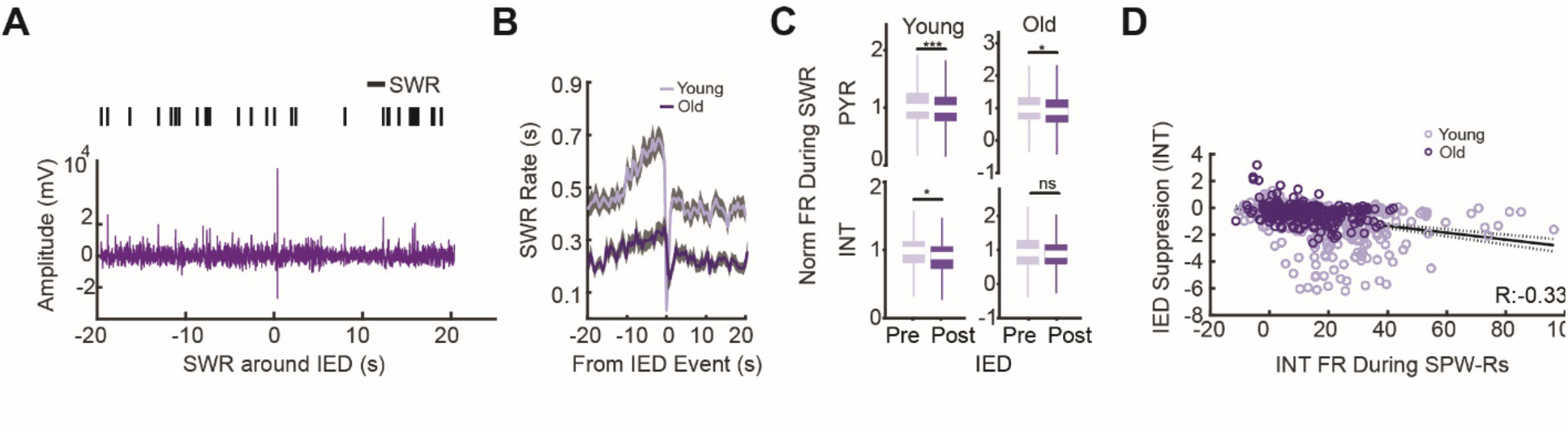
IEDs compete with SPW-R. **A**. Example trace of an IED (time 0) and SPW-Rs (black ticks) preceding and following the IED. **B**. Cross-correlograms between IEDs (time 0) and SPW-R occurrences in young (n = 43 sessions) and old (n = 36 sessions) mice. **C**. Group mean z-scored firing rates during SPW-Rs that occur– 10s and 0s before an IED (Pre) and 10s-20s after an IED (Post). (Wilcoxon rank-sum test, p = 9.77e-4, p=0.926; p=0.0217, p=0.9186; ns= not significant). **D**. Correlation between interneurons z-scored firing rate of interneurons post-IED and their z-scored firing rate during SPW-Rs (R=-0.33 p= 0.000). Light purple= young mice. Dark purple= old mice.

To compare the neuronal properties of IEDs and SPW-Rs associated neuronal activity, we plotted the peri-SPW-R spiking activity of neurons, sorted by the magnitude of their post-IED suppression. This comparison revealed that interneurons that were most effectively suppressed by IEDs corresponded to the neurons that were most effectively recruited during SPW-Rs (Fig.5D; Supplementary Fig. 4B-C).

### IEDs affect behaviorally relevant spike content of SPW-Rs

Since IEDs affected the incidence and associated firing rates of SPW-Rs, we next investigated whether spike content and waking experience-related replay of spike sequencies were affected by IEDs. Eight AD mice were trained on a figure-8 maze. As expected, pyramidal neurons had place fields (O’Keefe & Nadel, 1978) and the place fields tessellated the entire length of the maze (Fig. 6A). The place field sequences were “replayed’ in compressed sequences in either a forward or reversed manner during hippocampal SPW-Rs during sleep (Fig. 6; Davidson et al., 2009; Dragoi et al., 2011). The quality of replay, quantified by the trajectory score (see Methods), was similar between young and old animals (Supplementary Fig. 6). However, the fraction of significant replay events was lower in the 10 s epoch following IEDs compared to a similar epoch preceding them (Fig. 6C).

**Figure 6:**
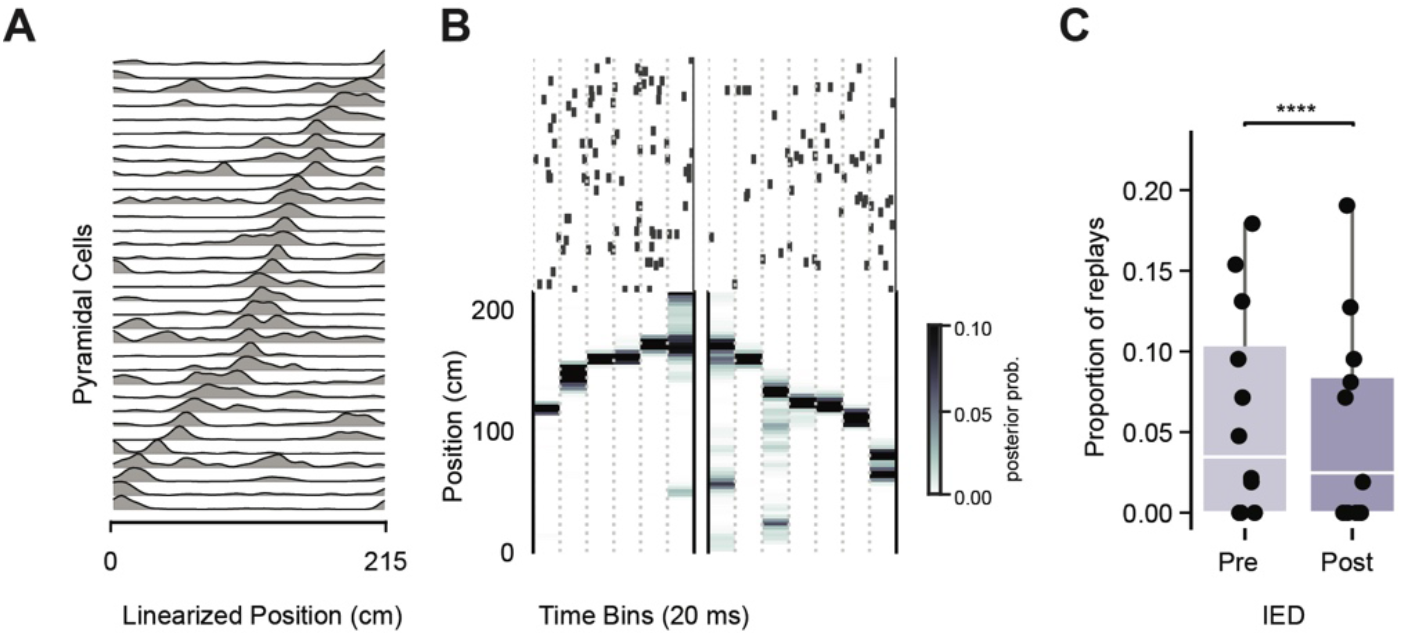
IEDs alter spike content of SPW-Rs. **A**. Place fields on a linearized T-maze track, sorted by the location of the peak firing rates. **B**. Example rasters of spikes (top) during SPW-R and the decoded virtual position of the animal (bottom) for forward and reverse replay events. Units in the raster (one neuron per row) were sorted according to the position of their place fields in the maze (A). **C**. Fraction of SPW-Rs with significant replay events during sleep that occurred in the –20 to –10s time window before IED (Pre) and in the 0 to 10s time window after IED (Post), P = 2.50 × 10−5, Generalized linear model.

### IEDs may affect memory performance

In contrast to behavior deficits seen in other AD mouse models of similar ages (Chen et al., 2000; Cayzac et al., 2015; Zhao et al., 2014; Jones et al., 2019), we did not find a significant difference between young and old APP/PS1 (C57BL/6;C3H) in the behavioral performance of the spontaneous alternation memory task (Supplementary Fig. 6A). However, the rate of IEDs (as quantified during comparable brain states of NREM sleep before the maze task), negatively and significantly correlated with daily memory performance in the figure-8 maze in old (R=-0.54, p= 0.0049) but not young (R=-0.23, p= 0.206) mice (Fig. 7A).

**Figure 7.**
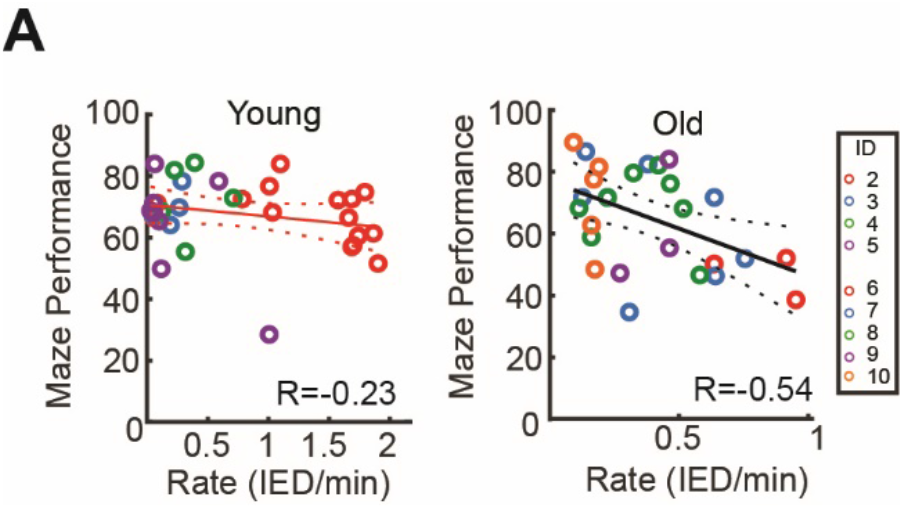
IEDs inversely correlate with memory performance in old AD mice. **A**. Correlation between IEDs rate in the home cage prior to daily maze performance and percentage of correct trials in the maze task for young (left) and old (right) mice (t-test, p=0.206; p=0.0049, respectively). Each color corresponds to a given animal.

## Discussion

Previous studies have reported altered physiological patterns in aged rodents and especially in AD mouse models (Radulescu et al., 2021; Palop & Mucke 2016; Wang et al., 2020; Mehak et al., 2022). AD pathology has been reported to correlate with impaired cognitive performance of the animals in various tasks (Lesné et al., 2008; Hsiao et al., 1996; Cramer et al., 2012; Zhang et al., 2011). The relationship between cognitive deterioration and physiological changes, however, is much less clear (Palop & Mucke 2016; Verret et al., 2012; Villette et al., 2010; Mehak et al., 2022; Wang et al., 2020; Bakker et al., 2012). It is possible that several overlapping and interrelated changes in neuronal dynamics contribute to decreased memory and other cognitive deficits. As the animal ages and pathology worsens, compensatory mechanism may play a role. Our work focused on the impact of IEDs on hippocampal dynamics, their relationship with SPW-Rs and memory. We find that IEDs in the AD mouse model are similar to IEDs in epileptic humans. IEDs were present at an early age in the AD mouse, in fact prior to histopathological changes and their incidence decreased with age (Reyes-Marin et al., 2017). IEDs reduce the probability of physiological SPW-Rs and altered their spike content. Furthermore, the incidence of IEDs inversely correlate with daily memory performance more in old rather than young AD mice. In depth examination of spiking activity of pyramidal cells and interneurons helped to resolve these apparently contradictory observations.

IEDs, by definition, refer to pathological population patterns of neurons between seizures (Alarcon et al., 1997; de Curtis et al., 2012). In experimental models, it has been assumed that seizures alter the excitability of a normal circuit and, in turn, give rise to transient events after the seizures (Alarcon et al., 1997; de Curtis et al., 2012; El-Hassar et al., 2007). Indeed, IEDs typically emerge after kindling (Goddard et al., 1969; Gelinas et al., 2016) and drug-induced seizures (Bragin et al., 2014; Curia et al., 2008). In contrast, we observed IEDs in APP/PS1 heterozygous male transgenic mice without any evidence of overt seizures. While it remains a remote possibility that rare seizures occurred sometime in the early life of the animal, a more likely possibility is that IEDs reflect pathological hyperexcitability due to various factors that may contribute to seizures or persist without inducing them. Thus, epileptiform discharge (ED) would be a more appropriate descriptor of these events and their development may not require *bona fide* seizures.

To support this hypothesis, we compared true IEDs recorded from the hippocampus in epileptic patients to IEDs in AD mice. The exact location of the tips of the wire electrodes could not be verified in patients. Nevertheless, large amplitude, negative, short duration (<10 ms) LFP events were recorded from wires which also recorded unit firing, an indication that the electrode was in the pyramidal layer. These putative population spikes were similar to events recorded in the CA1 pyramidal layer in rats (Buzsáki et al., 1991; Gelinas et al., 2016) and mice (Kam et al., 2016). IEDs were followed by a significant decrease of power in the high frequency oscillation band (80-160 Hz) in both humans and mice, and increased power in the theta (4–10 Hz) and gamma (30–70 Hz) bands, although these latter changes failed to reach significance in our epileptic subjects.

We separated putative pyramidal cells and interneurons in humans and found that both neuron types decreased their firing rates transiently after IED, again similar to observations in mice. Unfortunately, implanted patients, connected to recording equipment with inflexible cables, do not show normal sleep patterns. In fact, our patients were awake during most of the recording time in the afternoon and early night and, therefore, we could not determine the brain state-dependence of IED. In mice, the overwhelming majority of IEDs occurred during NREM sleep, in line with previous reports showing higher incidence of IEDs during NREM in human patients (Lambert et al., 2018). Overall, these similarities support our hypothesis that IEDs in AD mouse models are analogous to epileptic IEDs from patients.

IEDs occurred regularly in every recording session and in every AD mouse model tested to date (Verret et al., 2021; Reyes-Marin et al., 2017; Bakker et al., 2012; Kam et al., 2016; Brown et al., 2018). This contrasts with other rodent models, such as kainic acid model or pilocarpine model, which induce initial seizures or status epilepticus, followed by several weeks to months of incubation time before the emergence of IEDs and recurring seizures (Curia et al., 2008; Ben-Ari et al., 1978). The emergence of IEDs is hard to predict in these models, vary from subject to subject, and show large inter-subject variability (Leguia et al., 2021; Baud et al., 2018; Curia et al., 2008; Ben-Ari et al., 1978). Thus, AD mice might offer a better or alternative model for examining the molecular and physiological basis of IEDs and testing drug or other types of interventions for reducing their occurrence or improving their deleterious effects on other structures (Gelinas et al., 2016).

Long-term changes of IEDs and their impact on network dynamics were observed from the earliest testing age (2.5 months) of the AD mice. In agreement with other reports and in other AD mouse models, IEDs emerge prior to the documented molecular and histopathological changes (Busche et al., 2012; Kam et al., 2016; Reyes-Marin et al., 2017). Thus, IEDs are likely due to a hitherto unknown side effects of Aβ, the understanding of which would advance our knowledge of AD and the stability of network excitability (Busche et al., 2012; Zott et al., 2019; Walsh et al., 2002; Li et al., 2009 Xu et al., 2020; Palop & Mucke 2016; Verret et al., 2012).

IEDs not only emerged prior to AD pathology but their incidence and amplitude decreased over the lifetime of the animal (at least in the 2.5-13 months observation period; Reyes-Marin et al., 2017). One may interpret this as some kind of a compensatory mechanism that counters the hyperexcitability. As Aβ oligomers form plaques, there is switch to hypoactivity in AD mice (Busche & Konnerth, 2016). In-depth investigation of the neuronal mechanisms of IEDs in young and old mice suggests a potential mechanism. Average firing rates of pyramidal neurons, while showing wide, lognormal distributions (Mizuseki et al., 2011), were higher but less synchronous in old than in young AD mice, whereas firing rates of interneurons were lower in old mice. Depression of post-IED spiking activity of both pyramidal cells and interneurons was less pronounced in old mice at both transient (0.5 s) and sustained (20 s) intervals, even though the magnitude of synchronous discharge of neurons at the peak of the IED was comparable. Yet, the cross correlograms of all the neurons suggest that younger mice cells fired more synchronously than older neurons. Several seconds before IEDs interneurons ramped up or down their firing rates and this slope was steeper in young than in old mice. In young mice, the direction of firing change (ramping up or ramping down) was correlated with the neurons propensity to emit burst of spikes. In aged mice this correlation was absent. Overall, these observations demonstrate that as the animal ages, the overall E/I balance (Busche et al., 2012; Zott et al., 2019; Walsh et al., 2002; Li et al., 2009; Xu et al., 2020; Palop & Mucke 2016; Verret et al., 2012) is altered.

The altered E/I balance may also explain the relationship between IEDs and SPW-Rs. In young animals, the incidence of SPW-R increased steadily for several seconds prior to IEDs, presumably increasing, and potentiating the CA3-CA1 circuit to induce an IED, followed by a transient large decrease of SPW-R probability. This time course appeared to mimic the time course of firing rates of interneurons prior to IEDs. In contrast, the ramping of SPW-R incidence prior to IED was less pronounced in old AD mice, suggesting that less potentiation was required to reach the ‘threshold’ of IED occurrence.

IEDs may affect memory performance by altering SPW-Rs. It was hypothesized previously that IEDs my “hijack” SPW-Rs and alter their properties (Gelinas et al., 2016; Henin et al., 2022). We have also observed that these two hyperexcitable population events competed with each other. Increased SPW-R occurrence triggered IEDs and IEDs reduced the probability of occurrence of SPW-Rs. The qualitative features of SPW-Rs surrounding IEDs were also different. The probability of pyramidal cell and interneuron participation in SPW-R events was higher in pre-IED events and pre-post comparison was more pronounced in young mice.

Perhaps the most important feature contributed to SPW-Rs is their ability to replay neuronal sequences related to learned events in a compressed manner, assumed to be the mechanism of memory consolidation (Buzsáki et al., 1989; Wilson & McNaughton, 1994; Girardeau et al., 2009; Jadhav et al., 2012; Fernandez-Ruiz et al., 2019). The fraction of SPW-Rs with replay of place field sequences on the maze was reduced following IEDs. We also found a robust negative correlation between IED occurrence and memory performance in an alternation task and this inverse relationship was more robust in aged than in young mice. Mechanistic understanding of the complex relationship between IEDs, SPW-Rs, possibly other neurophysiological features, and memory impairment will need further clarification.

We also cannot ascertain whether the physiological differences we observed between young and old subjects reflect a normal aging process or AD pathology since WT mice did not have IEDs. Aged but non-AD rats show also show reduced incidence of SPW-Rs, reduced ripple frequency and altered spike coupling to various hippocampal patterns (Cowen et al., 2020; Crown et al. 2022; Wiegand et al., 2016), although it has not been examined how these changes might relate to behavior. A more extensive comparison of normally aged and AD mice may yield further clues. Another challenging task is to relate the evolving pathological changes to altered neuronal dynamic (Palop & Mucke et al., 2016) and testing whether artificial replication of the altered neuronal dynamic leads to similar behavior deficits.

## Acknowledgments

We would like to thank Andrea Navas Olivé, Annika Dhingra, Jiyun Shin, Anli Liu and the Experimental Pathology research core at NYU for their experimental support. This work was supported by NRSA 5TL1TR001447-07 (M.S.), FACES (M.S. and A.M.), ZKF Scientific Rotation Grant, Friedrich-Alexander University, Erlangen (A.M.) and NIH Grants MH122391 and U19 NS107616 (G.B.).

## Author Contributions Statement

M.S., A.M. and G.B. designed research; M.S., S.B., and H.H. performed research; M.S., A.M., M.V., R.H., and A.F.R. analyzed data; G.B. and M.S. wrote the paper.

## Methods

### Human Microwire recordings

We analyzed iEEG from 5 patients with drug-resistant epilepsy who underwent diagnostic depth electrode implantation for localization of the seizure onset zone. All patients were recruited in the preoperative epilepsy program of the Level 4 Epilepsy Center of Erlangen University Hospital, Germany. Indication for electrode implantation was made by the interdisciplinary epilepsy board. Ethical approval was granted by the “Ethik-Kommission” of Friedrich-Alexander Universität Erlangen-Nürnberg (142_12B). All patients gave informed written consent for the implantation of 1-2 hippocampal electrodes with microwire bundles. In addition to standard iEEG depth electrodes (Ad-Tech Medical, USA) implanted in temporal and extratemporal regions, in some patients 1-2 combined Macro/ Behnke-Fried Micro depth electrodes (Ad-Tech Medical, USA) were implanted in mesial temporal lobe structures. Electrode trajectories were planned by the clinical team based on surface EEG, seizure semiology and additional diagnostic modalities and were not influenced by this study. Electrode positions were confirmed by an intraoperative MRI- and postoperative MRI- and CT-scan. Only recordings from temporal electrodes with hippocampal microwires were included in this study. Microwire bundles consisted of 8 high-impedance wires and one low-impedance wire, which was used as reference. Following surgery, patients were monitored continuously for seven days upon withdrawal of antiepileptic drugs. Data was acquired using an ATLAS recording system consisting of CHET-10-A pre-amplifiers and a Digital Lynx NX amplifier (Neuralynx Inc, USA). Signals were filtered with cut-off frequencies at 0.1 and 9000 Hz and sampled at 4096 Hz for macro) and 32768 Hz for microwires.

Preprocessing: The raw data was converted to int16 dat format for processing with buzcode analysis scripts in Matlab (MathWorks; https://github.com/buzsakilab/buzcode). The LFP signal was downsampled to 2048 Hz. Spike sorting was performed semi-automatically and separately for each channel of Behnke-Fried Microwire signal using Kilosort (https://github.com/MouseLand/Kilosort), followed by manual curation of the waveform clusters with the Phy2 software (https://github.com/cortex-lab/phy). Neurons were characterized as pyramidal cells or interneurons according to their firing rate and waveform width in CellExplorer (Petersen et al., 2021). IED detection was performed on the microwire LFP signal from channels showing the highest number of pyramidal neurons firing in the respective patient.

### Experimental model and subject details

All experiments were approved by the Institutional Animal Care and Use Committee at New York University Langone Medical Center. Mice were kept in the vivarium on a 12-hour reverse light/ dark cycle in optimal ambient temperature (70°F) and humidity (50%) conditions. were housed two to three animals per cage before surgery and individually after surgery. Mice were provided food and water ad libitum. For behavior testing mice were water restricted to maintain 85% of their weight during and after behavioral training. We used littermates APP/PS1 heterozygous male transgenic mice (JAX Stock No. 004462) at ages 2-4 and 9-12 months.

### Surgical procedure

Mice were anaesthetized using isoflurane (1.5%–2%) in oxygen (30%) and placed in a stereotactic frame (David Kopf Instruments) as described previously (33). Craniotomies were made for either the hippocampus CA1 (AP:1.55 mm, ML:1 mm, DV:1.3 mm). H2 Silicon probes (Cambridge Neurotech) were mounted on custom-made 3D-printed drives and were inserted into the target brain region (Vöröslakos et al., 2021). Drives and a 3-D printed head cap were cemented to the skull with dental cement (C&B Metabond, #S380). Post implantation, craniotomies were covered with sterile wax. The copper-mesh surrounding the 3-D printed head cap was connected via a stainless-steel wire (A-M Systems, #792800) to a ground screw in the skull and served as a Faraday cage. Post-surgery, the probe was moved in the brain until the target region was physiologically identified.

### Recordings and Behavior

Electrophysiological recordings were conducted using an Intan RHD2000 interface board and 64-channel digital head stages with sampling at 20 kHz (IntanTech) and visualized using the Neurosuite software. The wideband signal was downsampled to 1250 Hz and used as the LFP signal. The Intan board was grounded, and a copper mesh placed under the recording cage was connected to the ground of the Intan board. Two weeks prior to behavioral experiments, animals were handled daily and accommodated to the experimenter and recording set up. Animals were water restricted before the start of the experiments. Mice were recorded at the same time every day for two hours in their home cage followed by 40 mins of running on a delayed (10 seconds) alternating figure 8 maze and ending with another two hours sleep recording in their home cage. During the behavioral task, the water deprived animal was rewarded when they alternated to the correct (left or right) corner. The animal’s position was monitored using a Basler camera (acA1300-60 gmNIR, Graftek Imaging) sampling at 30Hz to detect a head-mounted red LED. The automated maze was controlled by a custom-made Arduino-based circuit (https://github.com/valegarman).

### Unit clustering and neuron classification

Neuronal spikes were segregated using KiloSort (https://github.com/cortexlab/KiloSort) and manual curated using Phy (https://github.com/kwikteam/Phy). Putative excitatory and inhibitory neurons were separated based on their autocorrelograms, firing rates and waveform characteristics using CellExplorer (Petersen et al., 2021).

### Spectral analysis of LFP signals

Wavelet and spectral analysis of the LFP were analyzed throughout the experiment using routines written in MATLAB 2019 (MathWorks). The LFP was down sampled at 1250 Hz and low pass-filtered at 450 Hz. The power spectral density was computed using Welch’s average. The time-frequency spectrogram was computed between 2-100Hz using a moving window of 2 seconds. 60Hz (mice recording) and 50Hz noise (human recordings) was removed from all analysis.

### IED detection

The LFP signal (1250 Hz) was band-pass filtered between 20 and 80 Hz (Le Van Quyen et al., 2018) and transformed to a normalized squared signal (NSS). The standard deviation (SD) of the baseline homecage periods (excluding maze periods) was calculated. IEDs were defined by peaks beyond 10 (human) and 20 (mice) SD of the mean NSS whenever the signal crossed 3 (human) or 5 (mice) SDs of the mean NSS for 30 - 250 ms. All events were then visually inspected and manually curated. Whenever necessary the SD thresholds were adjusted for individual subjects. For further analysis, detected events were realigned to the peak of the unfiltered LFP. For mice, events from behavior periods were discarded because of artifact contamination. One-dimensional current-source density (CSD) signals were calculated using the second spatial derivative of LFP (Makarov et al., 2010; Mitzdorf 1985). Smoothing was applied to CSD signals for visualization purposes only. Tissue conductivity was considered isotropic (1 mV/mm2).

### Sharp-Wave ripple detection

Sharp wave ripple analysis was only done on epochs of sleep periods during the homecage recordings. Sleep epochs were identified by evaluating continuous LFP recordings in conjunction with EMG activity (Schomburg et al., 2014). Slow wave sleep (SWS) was characterized as <3Hz high amplitude LFP with low EMG. REM sleep episodes were identified by low EMG activity and high theta LFP power. While wake periods were identified by high EMG and theta LFP power. To detect sharp wave ripples, LFP signals from sites at the stratum pyramidal were bandpass filtered between 100-600 Hz as previously described (Tingley & Buzsáki, 2020). For ripple detection, the bandpass-filtered signal was subsequently smoothed using a Savitzky-Golay (polynomial) filter and candidate events were detected by thresholding (>2 SDs). Ripple duration limit were between 30ms to 100ms.

### Decoding/Replay

A Bayesian reconstruction approach was used to detect replay events (see (Johson & Redish 2007; Davidson et al., 2009; Dragoi et al., 2011; Grosmark & Buzsáki 2016; Pfeiffer & Foster 2015; Wu & Foster 2014; Zhang et al., 1998) for the use cases of this method for replay). Spike counts from each sharp wave ripple (SWR) were first binned into 20*ms* time bin *t* from *N* units *O*_*t*_ = (*o*_1*t*_*o*_2*t*_…*o*_*Nt*_) (200*ms* time bins were used for active behavior decoding). Then the posterior probability distributions for each *t* over binned positions (3 cm) along the linearized T-maze were calculated using Bayes’ rule.

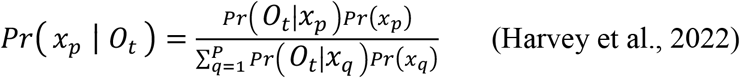

Where xp is the center of the p − th position bin. Assuming Poisson firing, the prior probability, Pr(Ot|xp,), for the firing of each unit n is equal to:

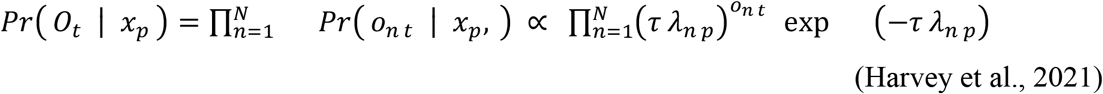

where *τ* is the duration of time (20 ms) and *λ*_*np*_ is the mean firing rate of the *n* − *th* unit in the *p*− *th* position. Assuming a uniform prior distribution *Pr*(*x*_*p*_) over the position bins. This implementation of Bayesian decoding was carried out using tools in the Python package Nelpy (Ackermann 2021). Each candidate replay event was considered a significant if the series of decoded positions were more consistent with an ordered trajectory compared to a surrogate distribution. Linear regression was used to fit a line to the posterior probability distribution. A Bayesian replay score for a given event was defined as the sum probability mass under the fit line within a bandwidth (21 cm) (Davidson et al., 2009). For each candidate event, we generated 1000 surrogates of the posterior probability distribution by circularly shifting each column of the posterior probability matrix by a random amount. A Monte Carlo p-value for each event was obtained from the number of surrogate events with replay scores higher than the observed score. The threshold for significance was held at 0.05. Replay events were identified from events with at least 5 active pyramidal cells with a peak firing rate of ≥ 1Hz and a peak-to-mean firing rate ratio of 1.5. In addition, candidate events with many (*>* 50%) inactive spike count bins were discarded. Finally, only sessions with *<* 25 cm median decoding error of the animal’s location during active running (speed *>* 3 cm/s) were used to quantify replay.

