## Supplementary Figures for "Interictal epileptiform discharges affect memory in an Alzheimer’s Disease mouse model"

**Table**

| | Patient Index | Recording Length (Hours) | Recording Start Time | Focus | Microelectrode position | Age | Gender |
|---|---|---|---|---|---|---|---|
| 1 | NY12 | 17.97 | 16:20 | T | R sub | 31 | M |
| 2 | NY13 | 15.04 | 17:48 | T | R DG | 20 | M |
| 3a | NY17 | 17.25 | 16:06 | F | R extrahippocampal | 30 | M |
| 3b | NY17 | 5.41 | 9:24 | F | R extrahippocampal | xx | xx |
| 4 | NY19 | 16.39 | 17:12 | T | L extrahippocampal | 52 | M |
| 5 | NY5 | 16.55 | 16:34 | T | R sub | 20 | M |
| 6 | NY6 | 16.84 | 15:29 | F | L DG | 26 | M |

**Supplementary Table 1: Clinical data from epilepsy patients included in the analysis.** Altogether 7 sessions from 6 patients were included (for patient 3, 3a and 3b are different recording sessions). Legend: Focus - seizure onset focus identified on intracranial EEG, T = temporal, F = frontal. Microelectrode position: position identified by co-registration of post-op CT and pre-op MRI (see methods). R = right, L = left , sub = subiculum, DG = dentate gyrus. M = male.

### Supplementary Figures

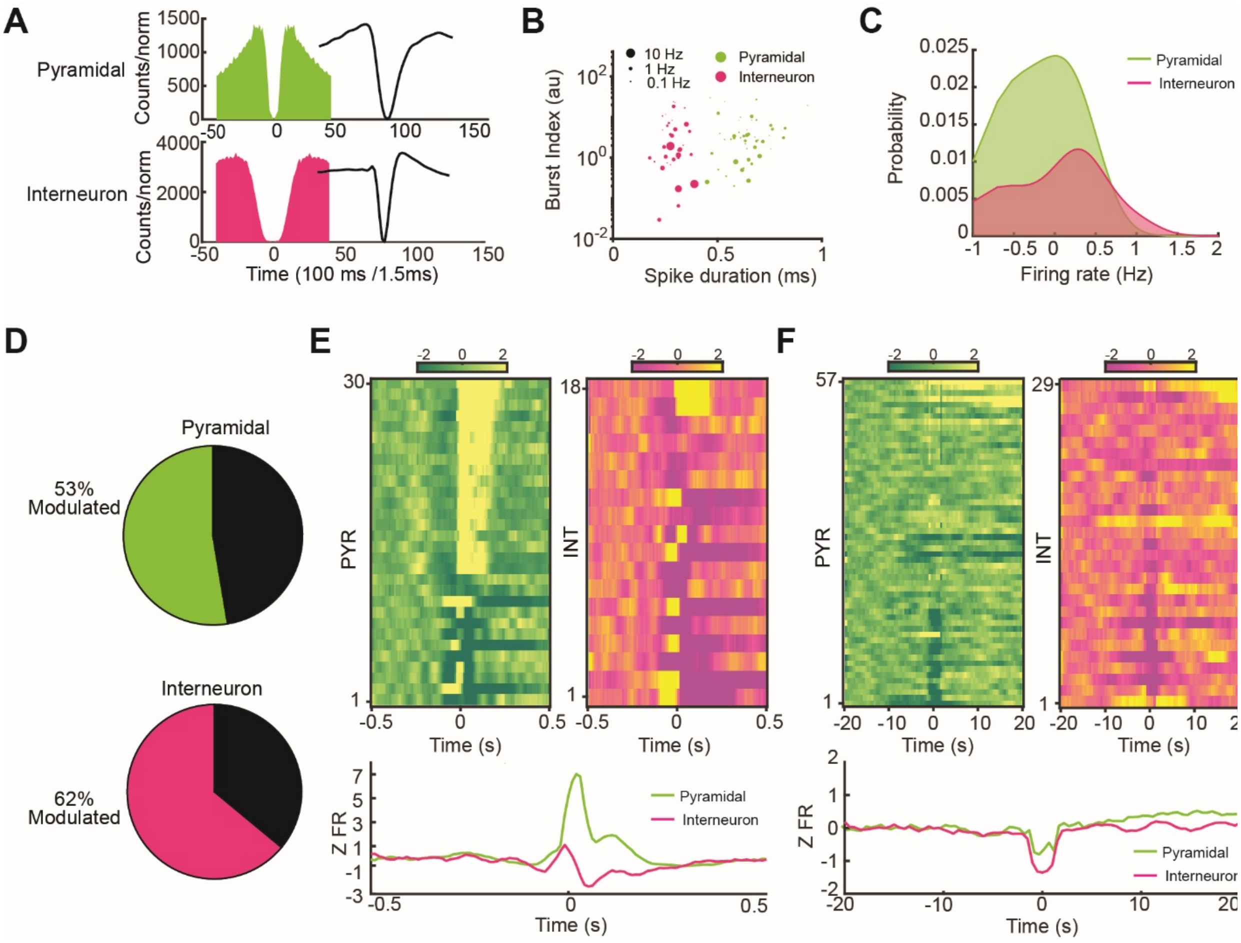


**Supplementary Figure 1: Properties of Human Unit Recordings**

**A.** Example auto-correlograms (left) and wide-band (1 Hz-20 kHz) action potential waveforms (right) of a human putative pyramidal cell and interneuron. **B.** Separation of putative pyramidal cells and interneurons on spike duration and bursting properties. Burst is defined as the fraction of spikes with < 6 ms. Spike duration: trough to peak latency. Each dot corresponds to one unit. Green dots are putative pyramidal cells and interneurons are in pink. Dot size represents relative firing rates. **C.** Probability distribution of log firing rates of pyramidal cells (green) and interneurons (pink). **D.** Percent of significantly modulated pyramidal cells (top) and interneurons (bottom) by IEDs. **E.** Top: Z-scored responses to IEDs of significantly modulated pyramidal cells (PYR) and interneurons (INT). Bottom: Average peri-IED response of modulated pyramidal cells (green) and interneurons (pink). **F.** Z-scored responses to IEDs of all pyramidal cells and interneurons 20s before and 20s after an IED. Bottom traces are the average z-scored response of all pyramidal cells and interneurons.

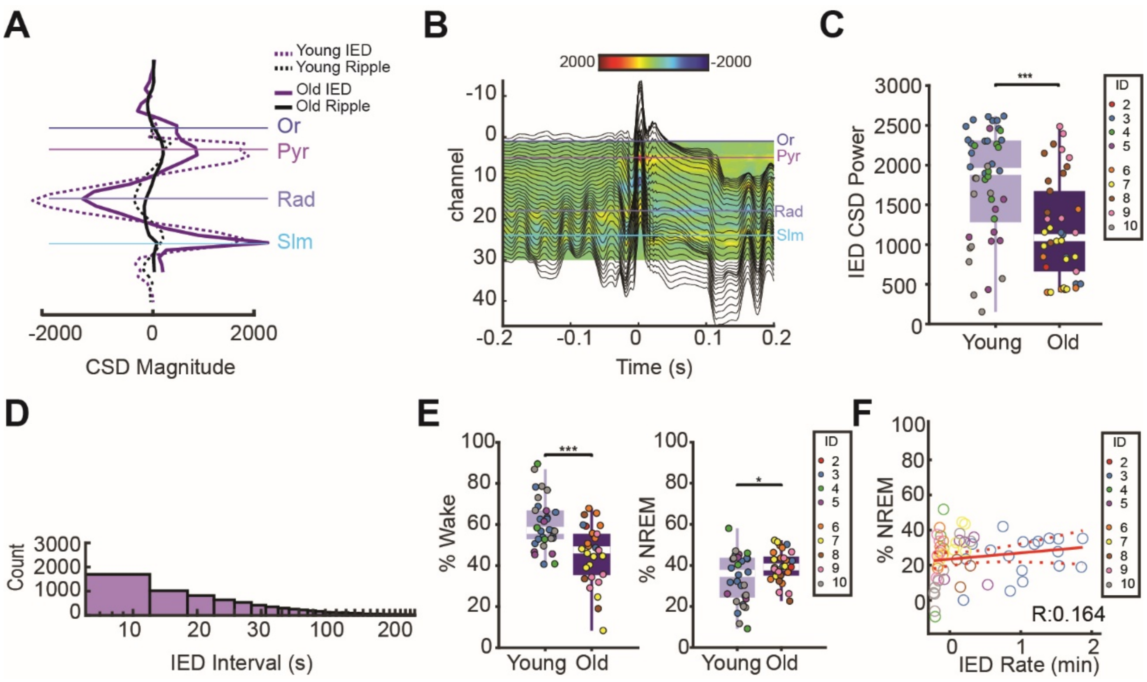


**Supplementary Figure 2: IEDs in APP/PS1 AD mouse**

**A.** Average CSD magnitude during IEDs and ripples in the same mouse at a young (3months; dashed lines) and old (9 months; solid lines) ages. Horizontal solid lines correspond to the str. oriens (Or), str. pyramidale (Pyr), str. radiatum (Rad), str. and lacunosum moleculare (Slm). **B.** Averaged LFP traces overlayed with the CSD map during an IED in an old mouse, complementing the IED CSD shown in Fig. 2C. **C.** IED CSD power between 0-0.05 s in young and old mice (two-sided Wilcoxon rank-sum test, p=1.9708e-4). **D.** Distribution of IED intervals (n =13,376from 83 sessions in 9 mice between all IEDs in seconds. X-axis is in log scale for visualization. **E.** Time spent in awake (right: two-sided Wilcoxon rank-sum test, p=5.13e-4) and in NREM (left: two-sided Wilcoxon rank-sum test, p=0.1433) in young and old mice. **F.** Correlation between IED rate and % of time spend in sleep per session. Each dot represents a session and each color, an animal (T-Test, p=0.2142).

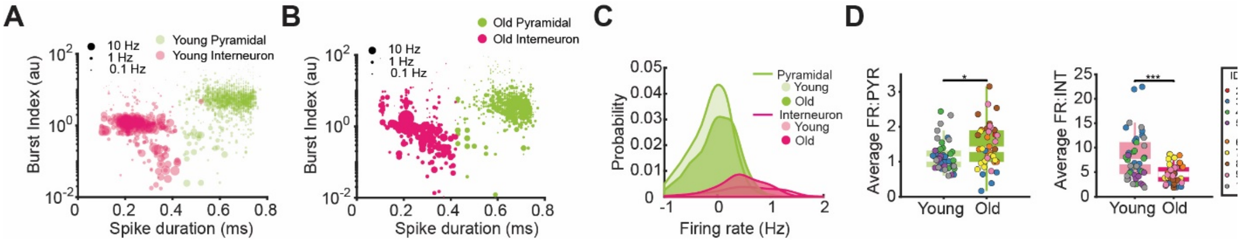


**Supplementary Figure 3: APP/PS1 AD mouse firing rate changes over time**

**A. B.** Separation of putative pyramidal cells and interneurons on spike duration and bursting properties in young and old AD mice. Burst is defined as the fraction of spikes with < 6 ms. Spike duration: trough to peak latency. Each dot corresponds to one unit. Green dots are putative pyramidal cells and interneurons are in pink. Dot size represents relative firing rates. **C.** Probability distribution of log firing rates for pyramidal cells (green:young- mean= -0.116 std=0.3975; old- mean= -0.002 std= 0.4094) and interneurons (pink: mean= -0.116 std=0.3975; old- mean= 0.4325 std=0.456) in young and old mice. **D.** Average log firing rates (y coordinate) of pyramidal cells (left: two-sided Wilcoxon rank-sum test, p= 0.017) and interneurons (right: two-sided Wilcoxon rank-sum test, p= 2.40e-4) in young and old mice.

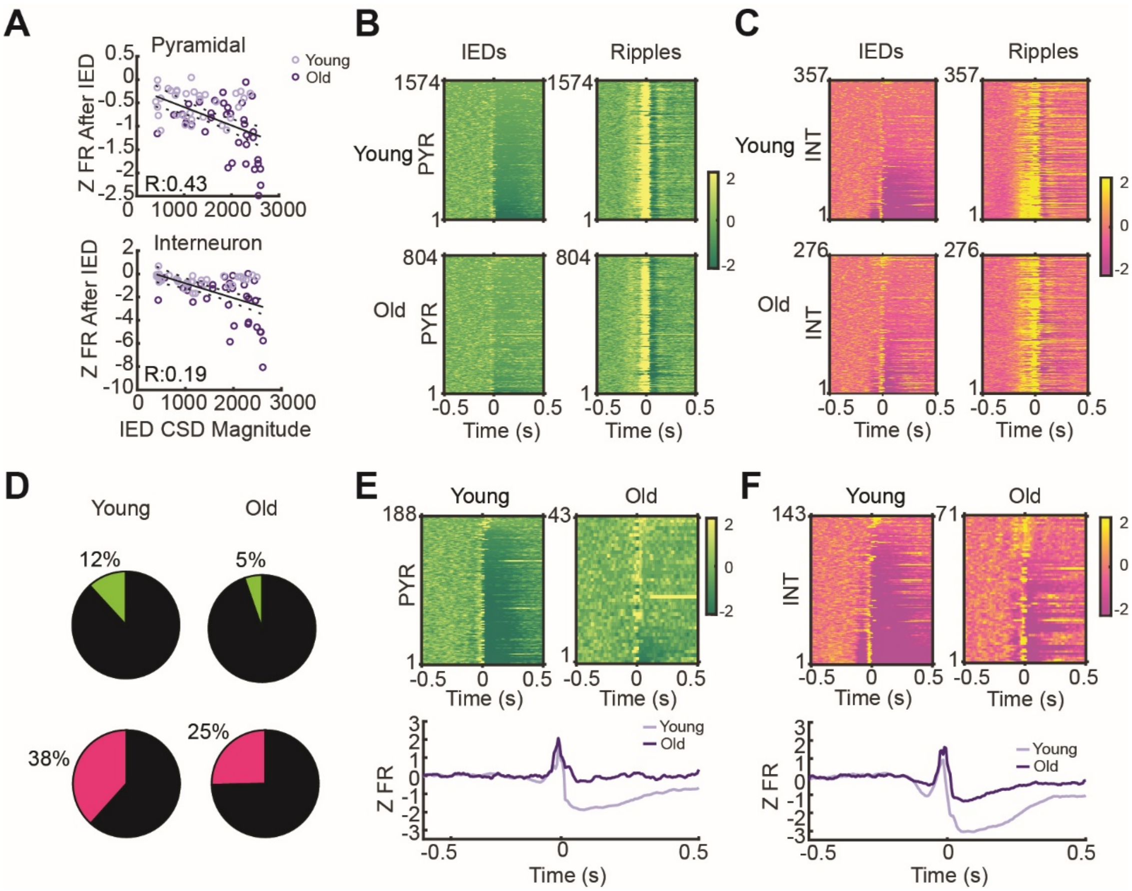


**Supplementary Figure 4: Spike correlations with IEDs in APP/PS1 AD mice.**

**A.** Correlation between Z scored firing rate and IED CSD magnitude in pyramidal cells (top: R=0.4304 p=5.432e-5) and interneurons (bottom: R=0.1907 p= 0.0882). Light purple= young mice. Dark purple= old mice. Sorted Z scored firing rates after IED in young and old pyramidal cells and interneurons as a function of the magnitude of IED CSD. Note that larger magnitude IEDs are followed by stronger suppression of spiking. **B.** z-scored peri-IED firing of pyramidal cells sorted by the strength of IED suppression (left). The same order of neurons is also shown for SPW-R-associated firing. **C.** Same as B for interneurons. All z-scored interneuron. **D.** Percent of significantly modulated pyramidal cells and interneurons by IEDs in young and old AD mice. **E. F**, In contrast to Fig. 4 A, B, these panels display peri-IED spiking only for the significantly modulated subsets of neurons. **E.** Left: Z-scored responses to IEDs of significantly modulated pyramidal cells. Bottom: Average IED-centered spiking of pyramidal cells for young and old AD mice. **F.** Same as E but for interneurons.

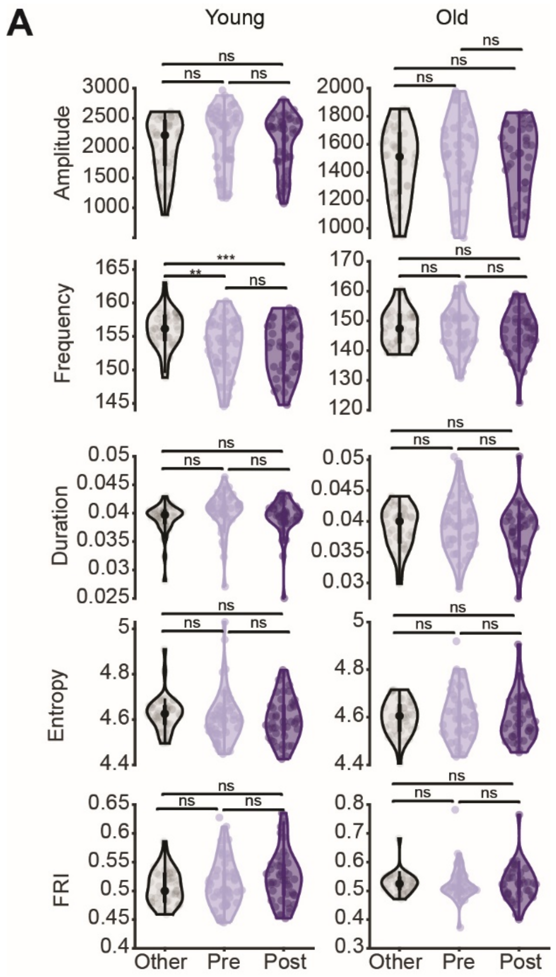


**Supplementary Figure 5: Ripple features around IEDs**

**A.** Group means of ripple amplitude (KW; young: p=0.334, p=0.779, p=0.7422. old: p=0.807, p=0.771, p=0.9978), frequency (KW; young: p=0.0255, p=0.0014 p=0.6444. old: p=0.814, p=0.321, p=0. 0.693), duration (KW; young: p= 0.013, p= 0.969, p=0.027. old: p=0.980, p=0.637, p= 0.760), entropy (KW; young: p= 0.222, p= 0.260, p= 0.995. old: p=0.927, p = 0.985, p= 0.857), and fast ripple index (FRI, KW; young: p=0.985, p= 0.285, p=0.373. old: p= 0.861, p= 0.969, p= 0.957) before and after IEDs, compared to ripples that occurred outside 40s (-20 to +20) around IEDs.  ns= not significant.

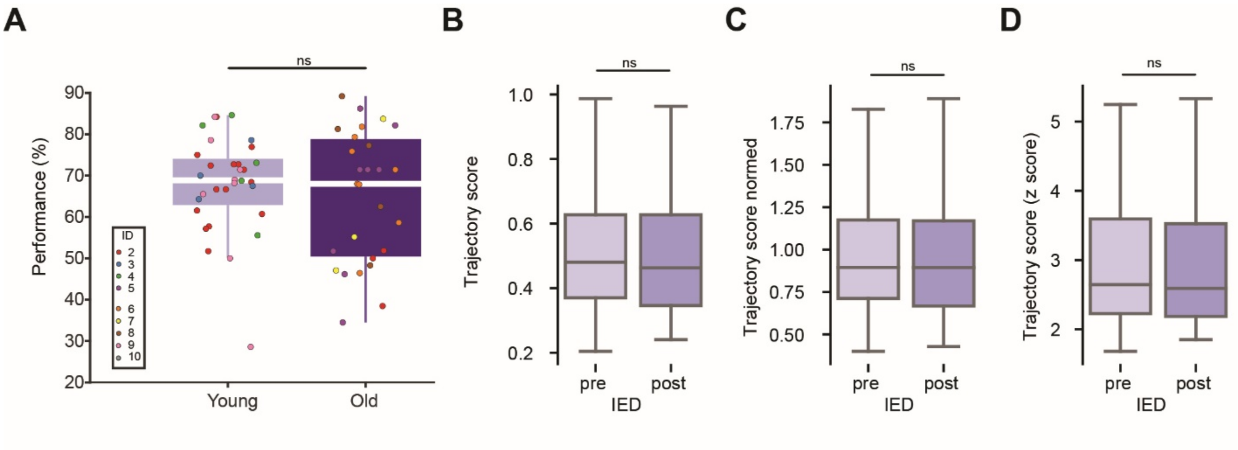


**Supplementary Figure 6: Replay quality of pre- and post-IED SPW-Rs is similar. A.** Average performance in young and old mice (KW, p= 0.842). Each dot corresponds to a session and each color to an animal. **B.** Trajectory score of replays during SPW-Rs before (–20s to –10s) and after (0 to 10a) IED (Mann-Whitney-Wilcoxon test two-sided, p = 0.6944). **C.** Trajectory score normalized by pre-sleep detected events (MannWhitney-Wilcoxon test two-sided, p = 0.5872). **D.** Z-scored trajectory score in units of standard deviations away from the Monte Carlo null distribution (Mann-Whitney-Wilcoxon test two-sided, p = 0.4414).

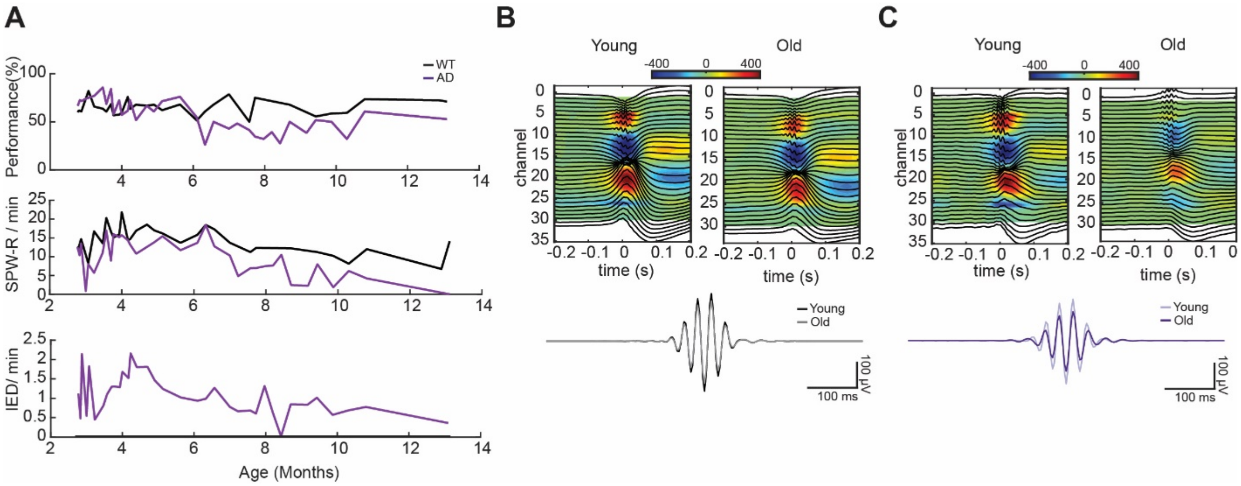


**Supplementary Figure 7: Chronic recordings are stable over 10 months. A.** Example performance, SPW-R rate and IED rate in 1 WT and 1 AD over 10 months. **B.** Average SPW-Rs in one session from the same WT animal in (A) at 3 months and 10 months. Bottom trace is an average of all the SPW-R in young mice (grey; 2-4 months) and old mice (black;9-12 months). **C.** Same as B but for AD mouse in (A).
